# Protective Population Behavior Change in Outbreaks of Emerging Infectious Disease

**DOI:** 10.1101/2020.01.27.921536

**Authors:** Evans K. Lodge, Annakate M. Schatz, John M. Drake

## Abstract

During outbreaks of emerging infections, the lack of effective drugs and vaccines increases reliance on non-pharmacologic public health interventions and behavior change to limit human-to-human transmission. Interventions that increase the speed with which infected individuals remove themselves from the susceptible population are paramount, particularly isolation and hospitalization. Ebola virus disease (EVD), Severe Acute Respiratory Syndrome (SARS), and Middle East Respiratory Syndrome (MERS) are zoonotic viruses that have caused significant recent outbreaks with sustained human-to-human transmission. This investigation quantified changing mean removal rates (MRR) and days from symptom onset to hospitalization (DSOH) of infected individuals from the population in seven different outbreaks of EVD, SARS, and MERS, to test for statistically significant differences in these metrics between outbreaks. We found that epidemic week and viral serial interval were correlated with the speed with which populations developed and maintained health behaviors in each outbreak.

## INTRODUCTION

One of the most important factors in assessing the danger posed by an epidemic of infectious disease is pathogen transmissibility. Widespread public anxiety during the 2013-2016 West African Ebola epidemic, while driven by an extremely high fatality rate during the early stages, was fueled in part by the speed with which Ebola Virus Disease (EVD) spread throughout the populations of Liberia, Sierra Leone, and Guinea.^1–3^ Similarly, concerns over annual influenza epidemics in the United States center on densely inhabited areas with multiple opportunities for viral transmission due to physical proximity between susceptible individuals.^4^ Regardless of geographic setting, understanding how to slow and control pathogen dissemination is a high priority in forecasting and preventing epidemics of infectious disease.

Epidemic modelers frequently employ compartmental models of disease outbreaks, such as Susceptible-Infected-Recovered (SIR) models as in Keeling and Rohani,^5^ Susceptible-Infected-Susceptible (SIS) models as in Gray *et al*,^6^ and Susceptible-Exposed-Infected-Recovered (SEIR) models as in LeGrand *et al.*^7^ Accurately estimating and modeling the number of infected and susceptible individuals in at-risk populations is of crucial importance in these models. Such estimation is complicated, however, by efforts to isolate infected individuals in hospitals or other settings to decrease contact with the susceptible population. While the isolation of infected individuals is beneficial and should be encouraged, it challenges data analysts because it is time-varying and reflects dynamic and often unpredictable human behavior. Moreover, the rate at which infected individuals are removed from the population typically accelerates throughout an epidemic as awareness of the infectious threat increases,^8^ a process Drake *et al* referred to as “societal learning.”^9^ Obtaining accurate estimates of this time-varying removal of infected persons, while difficult, improves the quality of compartmental models for epidemics of infectious disease.^9,10^ To our knowledge, however, no work has directly compared the rate of behavioral adaption across multiple epidemics, societies, and geographic settings.

Many factors can affect how quickly effective isolation practices are implemented, such as access to health care, local public health funding, international aid, and the efficacy of information campaigns.^11^ Local health care practices and non-formal healthcare systems also provide care to patients during epidemics and can play a part in quarantining infected individuals.^12^ Previous work in Liberia has shown that a combination of these approaches through simultaneous community engagement and clinical intervention is more effective than any single intervention, with both health care access and utilization increasing hand-in-hand to decrease EVD transmission during the 2013-2016 Ebola epidemic.^13^ While infection prevention often includes vaccination, progress to develop effective vaccines for emerging infections is slow and not necessarily more effective than isolation of infected individuals.^14^ Ring vaccination with the rVSV-ZEBOV-GP Ebola vaccine^15^ in the Democratic Republic of the Congo is promising,^16^ but previous work has suggested that ring vaccination may only provide a marginal benefit to rigorous contact tracing and patient isolation.^17^

The focus of this paper is the identification of key similarities and differences in the behavioral response to outbreaks of three emerging zoonotic infections. We sought to determine how the mean removal rate of infected individuals changed over the course of each outbreak as measured by epidemic week and viral serial interval. Individuals often experience zoonotic and emerging infections as innately more frightening than “familiar” diseases, leading to rapid behavioral adaptations due to high perceived risk.^18^ Behavior modification, while crucial for epidemic containment,^19–21^ is context dependent and difficult to predict due to social network, socioeconomic, and behavioral differences between populations.^22^ Thus, we chose seven different outbreaks of disease that stoked significant local and international fear due to the risk of global pandemic: the 2013-2016 Liberian Ebola epidemic, subsets of the 2013-2016 Liberian outbreak from Lofa and Montserrado Counties,^23^ the 2003 Hong Kong SARS epidemic,^24–26^ the 2014 Saudi Arabia MERS outbreaks in Riyadh and Jeddah,^27^ and the 2015 South Korea MERS outbreak.^28^ We examined whether epidemic week and serial interval successfully predicted days from disease onset to hospitalization (DSOH) and mean removal rate (MRR) throughout each epidemic.

## MATERIALS AND METHODS

### DATA

We obtained patient-level data for Ebola and MERS, and daily aggregated data for SARS. We added new columns to each epidemic dataset to track the number of days from symptom onset to hospitalization (calculated as hospitalization date - date of symptom onset; abbreviated DSOH) and the mean removal rate (calculated as 1 / DSOH; abbreviated MRR). In calculating MRR, we considered only positive DSOH values in order to focus on community transmission rather than nosocomial transmission. Additionally, we converted symptom onset dates to weekly onset dates by replacing each date with that of the closest previous Sunday.

### BINNED DATA

We compiled data for each outbreak location binned by epidemic week, to produce comparable data for regression analysis. Epidemic weeks came from weekly onset dates described above. We also binned the same data by serial interval, using 12 days as the estimated serial interval for Ebola,^23^ 8 days for SARS,^24^ and 7 days for MERS;^27^ this was calculated as epidemic week/(serial interval/7). Each dataset included, per week, the number of new cases, the cumulative number of cases, mean DSOH and associated standard deviation, and MRR and associated standard deviation. We removed epidemic weeks from the beginning of each outbreak so that the first three epidemic weeks had greater than 0 cases of disease each in order to focus on population-level behavioral adaptation to large-scale disease outbreaks instead of adaptations to individual disease events early in an epidemic. We performed all regression analyses using this binned data.

### REGRESSION ANALYSES

Initial regression analyses fit linear models to predict DSOH and MRR (Table 1, Eqs. 1-2). As before, data for DSOH excluded negative values (individuals who become symptomatic after being hospitalized for other reasons) to focus on community disease transmission and behavior change instead of nosocomial infection.

**Table 1.** Regression equations. DSOH stands for days from symptom onset to hospitalization; MRR stands for mean removal rate, calculated as (1 / DSOH). Epidemic weeks were weighted by cases per week. Outbreak location for DSOH and MRR included seven levels (Liberia, Lofa County, Montserrado County, South Korea, Riyadh, Jeddah, and Hong Kong).

| Eq. | Regression Type | Response | Predictor | Interaction Term |
| --- | --- | --- | --- | --- |
| 1 | linear | DSOH | epidemic week | none |
| 2 | linear | MRR | epidemic week | none |
| 3 | robust linear | DSOH | epidemic week | none |
| 4 | robust linear | MRR | epidemic week | none |
| 5 | robust linear | MRR | epidemic week | outbreak location |
| 6 | robust linear | MRR | serial interval | outbreak location |

Outlying points in the Liberian Ebola epidemic skewed our initial linear regression models. We compared manual removal of outliers, quantile regression, and robust linear regression to find the most appropriate method for handling such points. The three methods produced almost identical results. We used robust regression to re-fit all initial linear regression models to avoid the influence of outliers (Table 1, Eqs. 3-4). In addition, we performed robust linear regressions of MRR with an interaction term accounting for outbreak location (Table 1, Eq. 5-6) to examine predicted mean change in the MRR in each epidemic. We used the Bonferroni correction^29^ for multiple comparisons to compute confidence intervals, utilizing a 99% confidence interval in our model comparisons. The size of the smaller epidemics (MERS and SARS) played a large part in determining confidence interval size and significance. All data management, modeling, and visualization was performed in R.^30^

## RESULTS

### DAYS TO HOSPITALIZATION (DSOH) AND MEAN REMOVAL RATE (MRR)

DSOH consistently declined over time in each epidemic. Robust regressions for DSOH and MRR (Table 1, Eqs. 3 and 4) showed negative and positive slopes, respectively, which corroborated the observations made on non-binned data (Fig. 1).

**Figure 1.**
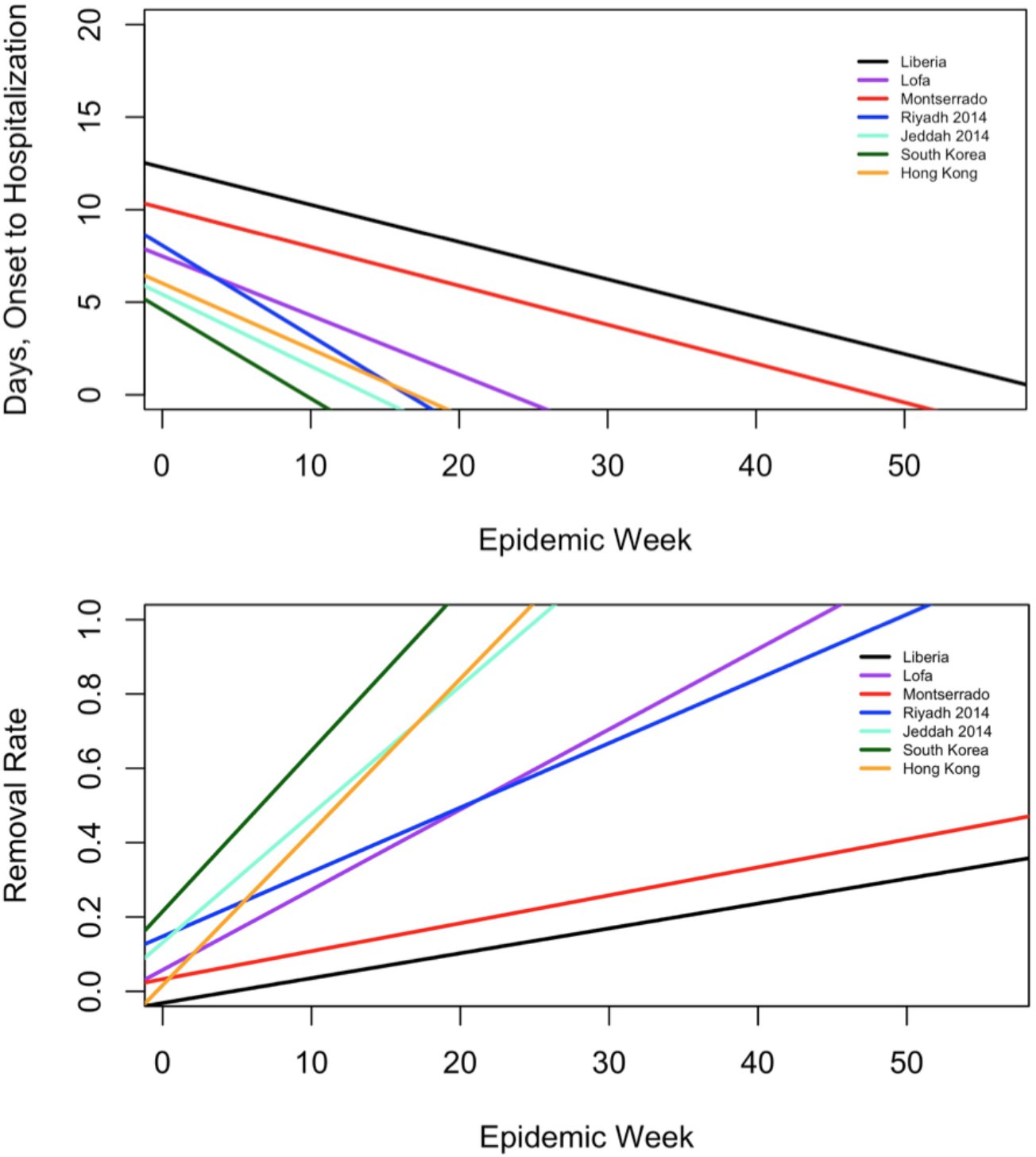
Regression: DSOH and MRR. The public health response to epidemic infection varied widely between the outbreaks studied. These graphs depict model lines from regressions of each of the 2 response variables (DSOH (Table 1, Eq. 4) and MRR (Table 1, Eq. 5)) on epidemic week for the 7 outbreaks indicated in the legend. South Korea and Liberia exhibited the most extreme slopes in both analyses. As an illustration of the observed difference between outbreaks, the graphs show South Korea achieving an almost complete removal of infected individuals from the population and a sharp decline in days till hospitalization within 20 weeks, while Liberia only achieved a roughly 20% removal rate by 50 weeks.

From robust regression analyses accounting for outbreak location (Table 1, Eqs. 5 and 6), we calculated the mean change in the MRR for each outbreak location using the *interactionMeans* function from the R package phia for post-hoc interaction analysis. This analysis showed that the mean change in the MRR of the Hong Kong SARS epidemic was approximately five times (per serial interval) to seven times (per epidemic week) more than the mean change in the MRR of the Liberian Ebola epidemic (Fig. 2). The mean change of the MRR in the Ebola epidemic in Lofa County, Liberia, was significantly higher than the mean change of the MRR for the overall Liberian epidemic and the outbreak in Montserrado County, Liberia, regardless of predictor (epidemic week or serial interval) (Fig. 2). The three MERS outbreaks (Riyadh, Jeddah, and South Korea) did not differ significantly from one another and had limited precision (Fig. 2).

**Figure 2.**
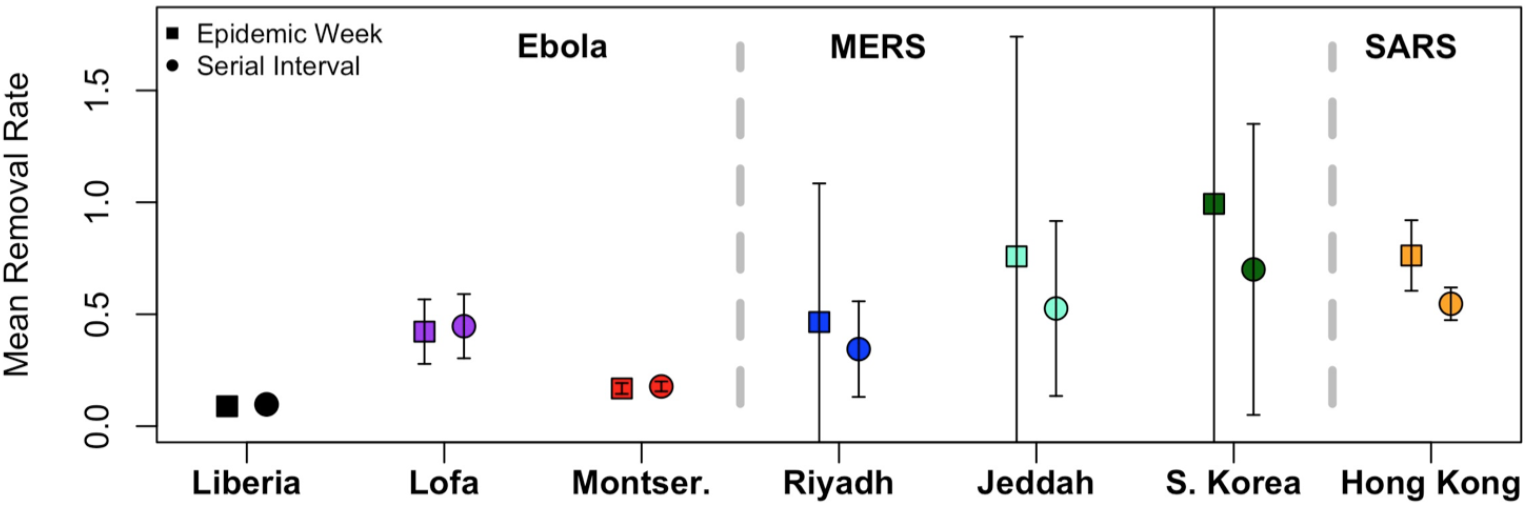
Adjusted MRR. The average change in the mean removal rate (MRR) differed based on the type of outbreak, the location, and the independent variable (epidemic week or serial interval) used to predict changes in MRR. The error bars plotted around the estimated MRR values represent the normalized 99% confidence intervals calculated using the *interactionMeans* function in R. The most precise estimates were observed for the Ebola outbreak, and the largest for the outbreaks of MERS. Predicting the mean change of the MRR with serial interval generally led to tighter confidence intervals than predictions using epidemic week. These results indicate that an outbreak’s type and location are important determinants of the mean change in the MRR per epidemic week or serial interval.

We found that predicting mean change of the MRR by epidemic week (Table 1, Eq. 5) led to higher mean estimates and wider confidence intervals in the MERS and SARS outbreaks; conversely, predicting with serial interval (Table 1, Eq. 6) lowered mean estimates and narrowed the associated confidence intervals (Fig. 2). We identified little difference in the mean change of the MRR for the Ebola outbreak depending on predictor (Fig. 2). This indicates that, at least in the case of MERS and SARS, both the passage of time and the serial interval of each virus may affect the speed with which populations develop and maintain health behaviors.

## DISCUSSION

The primary finding of this study was that removal of infected individuals from the susceptible population, measured as DSOH and MRR, increases over time and varies significantly based on outbreak duration and location. While DSOH improved (decreased) in every epidemic over time, extreme disparities in starting values (approximately 13 days from symptom onset to hospitalization at the beginning of the 2013-2016 Ebola outbreak in Liberia, versus approximately 5 days in the 2015 MERS outbreak in South Korea) highlight the intrinsic disadvantage that low-income countries may experience due to the interrelated concerns of poverty, limited access to health care, and low investment in public health. DSOH and MRR regressed against epidemic week differed across all observed outbreaks, and MRR likewise differed markedly based on the virus in question (Ebola, MERS, or SARS), the location, and at times both. Both DSOH and MRR are useful measurements of public health behavior during outbreaks, and are useful tools to compare outbreak response effectiveness in distinct geographic, economic, and social settings. Of course, DSOH and MRR are intrinsically and simply related since one is simply the reciprocal of the other. The main advantage of DSOH is that it is expressed in intuitive units (days elapsed), whereas MRR reflects the theoretical “removal rate” of standard compartmental models.^5^

Figure 2 highlights differences in mean change of the MRR due to outbreak type (Ebola, MERS, or SARS) and location. Mean change of the MRR was similar when calculated using epidemic week versus serial interval for Ebola, but demonstrated a lower estimate and lower standard error when calculated using serial interval in all three outbreaks of MERS and the outbreak of SARS in Hong Kong. This suggests that the relevance of various predictors (epidemic week versus serial interval) may vary based upon the type and location of an outbreak, although the comparative relevance of epidemic type versus location cannot be disentangled with the data available in this study. We recommend similar analyses of MRR be conducted across a wide range of geographies as outbreaks of emerging pathogens arise, providing important data on the range of MRR, and its expected rate of change, in different settings.

While our findings demonstrate large and statistically significant differences in MRR, it is notable that the calculated rates of change in the MRRs are within a factor of ten (when calculated using epidemic week) to seven (when calculated using serial interval) of each other (Fig. 2), with the mean change being the lowest in the EVD outbreak in Liberia and the highest in the MERS outbreak in South Korea. For modelers seeking to understand the epidemiology of emerging infectious diseases with limited or no data from previous outbreaks, this study provides a range of acceptable values for the MRR based on seven geographically distinct outbreaks of three emerging diseases. Similarly, while large disparities in DSOH are obvious (Fig. 1), these data highlight that all societies quickly adapt to outbreaks of emerging infections. Drake *et al* previously demonstrated the positive impact of behavior change in infectious outbreaks, noting that doubling the rate of “societal learning” in a model of the 2003 SARS outbreak in Singapore approximately halved the estimated number of infected patients.^9^ While there is a theoretical upper limit to the speed with which newly-infected individuals can be removed from the susceptible population,^9^ public health strategies aimed at fostering behavioral adaptations and accelerating isolation should form a cornerstone of interventions tasked with limiting the spread of highly contagious and deadly emerging pathogens.^31^

## CONCLUSION

We have shown that public health practices for isolating infected individuals from the susceptible population vary significantly by pathogen and location, but can in some cases be predicted by the timing and serial interval of the epidemic. This study detected variation in DSOH and MRR based on epidemic location and outbreak type, indicating that it may be possible to estimate a general range of the rate of change in these variables over time. Due to location-specific differences in DSOH and MRR, modelers who seek to develop forecasts early in an outbreak would benefit from estimating an expected range for removal of infected individuals using data from past outbreaks of the same pathogen in a similar setting. Furthermore, the quality of these estimates will be impacted by the metric chosen, as seen by the notable, but distinct, trends detected in DSOH and MRR. As seen in this study, utilizing a well-chosen response variable with a relatively small amount of data can provide material for making effective forecasts about public health behavior.

## DATA, CODE, AND MATERIALS

We studied seven outbreaks: the 2013-2016 Liberian Ebola epidemic on a country-wide level, subsets of the same epidemic in Lofa and Montserrado Counties, the 2003 Hong Kong SARS epidemic, the 2014 Saudi Arabia MERS outbreaks in Riyadh and Jeddah, and the 2015 South Korea MERS outbreak. The Ebola data was originally obtained by the World Health Organization and provided by Christopher Dye. The Hong Kong SARS data was provided by Gabriel Leung of Hong Kong University. Please contact Christopher and Gabriel for data regarding Ebola and SARS, respectively, due to concerns regarding potentially identifiable health information. Finally, the MERS data for Saudi Arabia and South Korea were obtained from data compiled by Andrew Rambaut of the University of Edinburgh, and is publicly available at https://github.com/rambaut/MERS-Cases/blob/gh-pages/data/cases.csv.

## AUTHOR’S CONTRIBUTIONS

Evans Lodge helped develop the project and conduct initial data analysis with John Drake as part of the University of Georgia Population Biology of Infectious Diseases REU Site. Annakate Schatz updated Lodge’s analyses with finalized Ebola data and wrote an initial paper draft from that work. Lodge and Drake finalized the manuscript for submission.

